# IL4Rα Controls The Medullary Dwell Time Of Mature Thymocytes By Scheduling Their Release From The Thymus

**DOI:** 10.64898/2026.11.07.765082

**Authors:** Kieran D. James, Andrea J. White, Fabrina M. Gaspal, Sonia M. Parnell, Andrea Bacon, Stefano A. P. Colombo, Andrew S. MacDonald, Anne Wolf, Maria Milani, Thomas Langmann, William E. Jenkinson, Graham Anderson

## Abstract

Following their generation, newly produced single positive (SP) thymocytes must remain in the thymus to undergo post-selection maturation, a process supported by the thymic medulla. This ensures that SP thymocytes are equipped with functional competence prior to S1PR1-mediated thymic emigration. Despite this importance, the intrathymic mechanisms that control the balance between retention and release of SP thymocytes is unclear. Here, we describe a cell autonomous requirement for IL4Rα-signalling in SP thymocytes, that determines the duration of their medullary occupancy. We show that the period of this occupancy is prolonged in the absence of IL4Rα expression by SP thymocytes, which occurs as a result of impaired interactions with medullary thymic epithelial cells (mTEC). Importantly, extending the duration of medullary occupancy enhances the maturational status and functional capabilities of SP thymocytes, including improved TNFα and IL2 production. Collectively, we show that IL4Rα regulates the medullary occupancy of mature thymocytes, which in turn determines the functional properties of thymic output.

## Introduction

During αβT-cell development in the thymus, immature CD4^+^CD8^+^ and CD4^+^8^−^/CD4^−^8^+^ thymocyte subsets undergo a series of positive and negative selection events that determine whether they die intrathymically or become mature T-cells^1–3^. The outcome of this thymic selection process depends upon the ability of randomly generated αβT-cell receptors (αβTCRs) to recognise self-peptide/MHC complexes expressed by cells of the thymic microenvironment, including cortical thymic epithelial cells (cTEC), medullary thymic epithelial cells (mTEC) and dendritic cells (DC)^4,5^. Successful completion of thymic selection results in the generation of self-tolerant CD4^+^ and CD8^+^ single positive (SP) thymocytes that reside within the thymic medulla and are restricted to MHC II and MHC I respectively. SP thymocytes are then released from the thymus into the periphery, a process known as thymic emigration or egress, where they are termed recent thymic emigrants (RTE)^6,7^. These RTE contribute to the maintenance of a diverse naïve T-cell pool that mediates effective immune responses to pathogens.

Although the thymic medulla acts as a repository for SP thymocytes, not all SP thymocytes are the same. Early studies showed that SP4 thymocytes consist of both functionally immature and mature subsets^8^, which provided strong evidence for the need for post-selection intrathymic differentiation events that enable newly produced T-cells to acquire functional maturity. More recent studies have shown this heterogeneity, and the need for post-selection maturation, can be mapped to defined SP subsets identified using various markers including MHC I, CD69 and CD62L^9,10^. For example, while CD69^+^MHC I^−^ semi mature (SM) SP thymocytes are functionally immature, downstream mature 1 (M1) CD69^+^MHC I^+^ thymocytes undergo proliferation in response to TCR signalling, while the most mature 2 (M2) CD69^−^MHC I^+^ SP thymocytes can also produce cytokines following TCR ligation, a process termed cytokine licensing^9^. M2 SP4 thymocytes are also selectively equipped with expression of S1PR1, a cell surface receptor that enables the egress of mature SP thymocytes from the thymus^11–13^. In order for these multiple events to occur during post-selection maturation, SP thymocytes are retained in the thymus for 4-5 days following their generation^14,15^. Importantly, and despite being a critical step in functional T-cell production, the mechanisms that determine the duration of this intrathymic dwell time period are poorly understood.

Here, we show IL4 receptor α (IL4Rα) signalling in SP thymocytes controls the duration of their medullary occupancy in a cell autonomous fashion. Mechanistically, selective deletion of IL4Rα in SP thymocytes was found to prolong their intrathymic dwell time as a result of impaired interactions with medullary thymic epithelial cells (mTEC). Importantly, IL4Rα-deficient thymocytes can exploit this extended thymus occupancy by continuing their post-selection maturation, which results in enhancement of their functional properties. Collectively, we describe the importance of IL4Rα in late-stage thymocyte development, and demonstrate that post-selection maturation is a continuous process determined by time and not developmental stage.

## Results

Analysis of Thymocyte Development Following CD4^Cre^ Mediated Deletion of IL4Rα. In recent years, several studies have demonstrated the presence of multiple components of type 2 immune networks in the thymus^16–18^. Of these, the type 2 cytokine IL4, which binds to the γc/IL4Rα complex on thymocytes, is readily detectable intrathymically at steady-state^19,20^. To examine a possible role for IL4/IL4Rα in the regulation of thymocyte development, we first analysed IL4Rα expression at specific stages of thymocyte development. Flow cytometric analysis with anti-IL4Rα antibodies detected expression in CD4^+^CD8^+^ (DP) and downstream conventional CD4SP (cSP4, CD4^+^CD8^−^TCRβ^hi^CD25^−^PBS57-CD1d tetramer^−^), at each maturational stage (CD69^+^MHC I^−^ SM, CD69^+^MHC I^+^ M1 and CD69^−^MHC I^+^ M2) (Fig. 1A). Consistent with this, *in vitro* stimulation of WT thymocytes with recombinant IL4 (rIL4) induced robust expression of phosphorylated STAT6 (pSTAT6), a downstream indicator of IL4Rα signalling (Fig. 1B)^19,21^. Given the broad expression of IL4Rα by multiple haemopoietic and non-haemopoietic cell types, and to specifically analyse its role in later stages of thymocyte development, we crossed *Cd4^Cre^* and *Il4ra^fl/fl^* mice to generate *Cd4^Cre^Il4r^fl/fl^* mice, where the *Il4ra* subunit is conditionally deleted in DP and SP thymocytes. Successful and efficient deletion was confirmed by demonstrating a loss of pSTAT6 signalling in both DP and SP thymocytes of *Cd4^Cre^Il4ra^fl/fl^* thymocytes (Fig. 1B), and absence of detectable IL4Rα expression following staining of *Cd4^Cre^Il4ra^fl/fl^* thymocytes with an anti-IL4Rα antibody (Fig. 1C). Initial analysis of the thymus in *Cd4^Cre^Il4ra^fl/fl^*mice and *Cd4^Cre^* controls showed comparable total cellularity (Fig. 1D), with no gross differences in major thymocyte subsets, including DP, total cSP4, and total TCRβ^hi^ SP8 cell numbers (Fig. 1D). When thymic sections of *Cd4^Cre^* and *Cd4^Cre^Il4ra^fl/fl^*were stained with antibodies to CD4 and CD8 alongside the medullary thymic epithelial cell (mTEC) marker ER-TR5, confocal analysis showed the presence of organised cortical and medullary ares that contained DP and SP thymocytes (Fig. 1E). Collectively, these data indicate that thymocyte expression of IL4Rα is not required for the generation of SP thymocytes from DP progenitors, nor the positioning of these cells in their relevant thymic microenvironments.

**Figure. 1:**
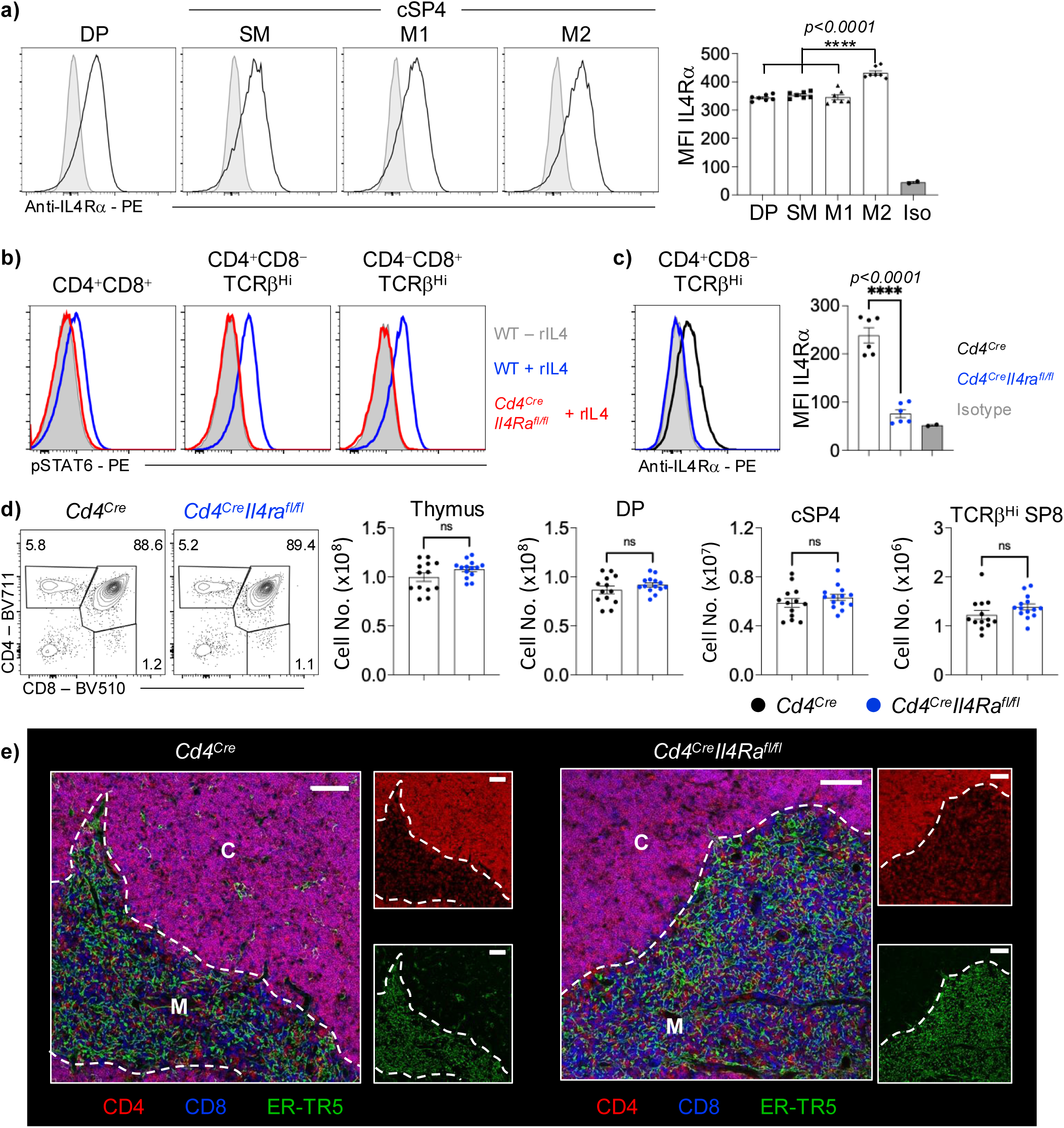
Late Stages of Thymocyte Development Occur Independently of *Il4ra.* **(A)** Flow cytometric analysis of IL4Rα expression by CD4^+^CD8^+^ (DP) thymocytes, and SM, M1 and M2 conventional SP4 (cSP4; PBS57-tetramer^−^CD4^+^CD8^−^ TCRβ^hi^CD25^−^) thymocytes. Mean fluorescence intensity (MFI) for each population is plotted on the right-hand side. Plots representative of n=7 from 2 independent experiments. **(B)** Flow cytometric analysis of pSTAT6 expression by unsorted DP, CD4^+^CD8^−^TCRβ^Hi^ and CD4^−^CD8^+^TCRβ^Hi^ thymocytes from WT or *Cd4^Cre^Il4ra^fl/fl^*, following 10-minutes incubation in the presence or absence of *10ng/ml* rIL4. Histogram plots are representative of n>4 from 2 independent experiments. **(C)** Flow cytometric analysis of IL4Rα expression by CD4^+^CD8^−^TCRβ^Hi^ from *Cd4^Cre^* (Black) and *Cd4^Cre^Il4ra^fl/fl^* (Blue), alongside an isotype control (Grey). Plots representative of n=6 from 2 independent experiments. **(D)** Flow cytometric analysis of CD4 and CD8 expression by thymocytes from *Cd4^Cre^* (left, Black) and *Cd4^Cre^Il4ra^fl/fl^* (right, Blue) adult mice. Absolute numbers were calculated for thymic cellularity, DP, cSP4 and CD4^−^CD8^+^TCRβ^Hi^ SP8 thymocytes. Plots representative of n=13 for *Cd4^Cre^* and n=14 for *Cd4^Cre^Il4ra^fl/fl^* from 6 independent experiments. **(E)** Confocal microscopy of thymus sections from *Cd4^Cre^* (left) and *Cd4^Cre^Il4ra^fl/fl^* (right) stained for CD4 (Red), CD8 (Blue) and ER-TR5 (Green) to determine thymocyte distribution. Scale bar indicates (100μm), images are representative of >3 images each from 3 *Cd4^Cre^* and 3 *Cd4^Cre^Il4ra^fl/fl^*, from total of 2 independent experiment.

### Thymocyte Expression of IL4Rα Determines the Intrathymic Dwell Time of Mature Thymocytes

To further examine the impact of IL4Rα deficiency on post-selection thymocyte maturation, we crossed *Cd4^Cre^Il4ra^fl/fl^* mice with Rag2GFP reporter mice^7^, and sub-divided bulk cSP4 thymocytes into semi-mature (SM), mature 1 (M1) and mature 2 (M2) on the basis of their expression of CD69 and MHC I^9^ (Fig. 2A, B; Fig. 2 Supp Fig. 1A). Importantly, such analysis demonstrated a stage-specific increase at the Rag2GFP^+^ CD69^−^MHC I^+^ M2-stage of cSP4 maturation (Fig. 2B). As expression of GFP identifies new thymocyte production, this rules out the possibility that increased SP thymocytes are a result of increased recirculation of Rag2GFP^−^ peripheral T-cells back to the thymus. Furthermore, when we analysed the proliferative status of thymocytes, we saw comparable staining for Ki67 in DP and cSP4 M2 thymocytes from *Cd4^Cre^*and *Cd4^Cre^Il4ra^fl/fl^*Rag2GFP mice (Fig. 2C), indicating that increased SP frequency was not due to their increased proliferation.

**Figure. 2:**
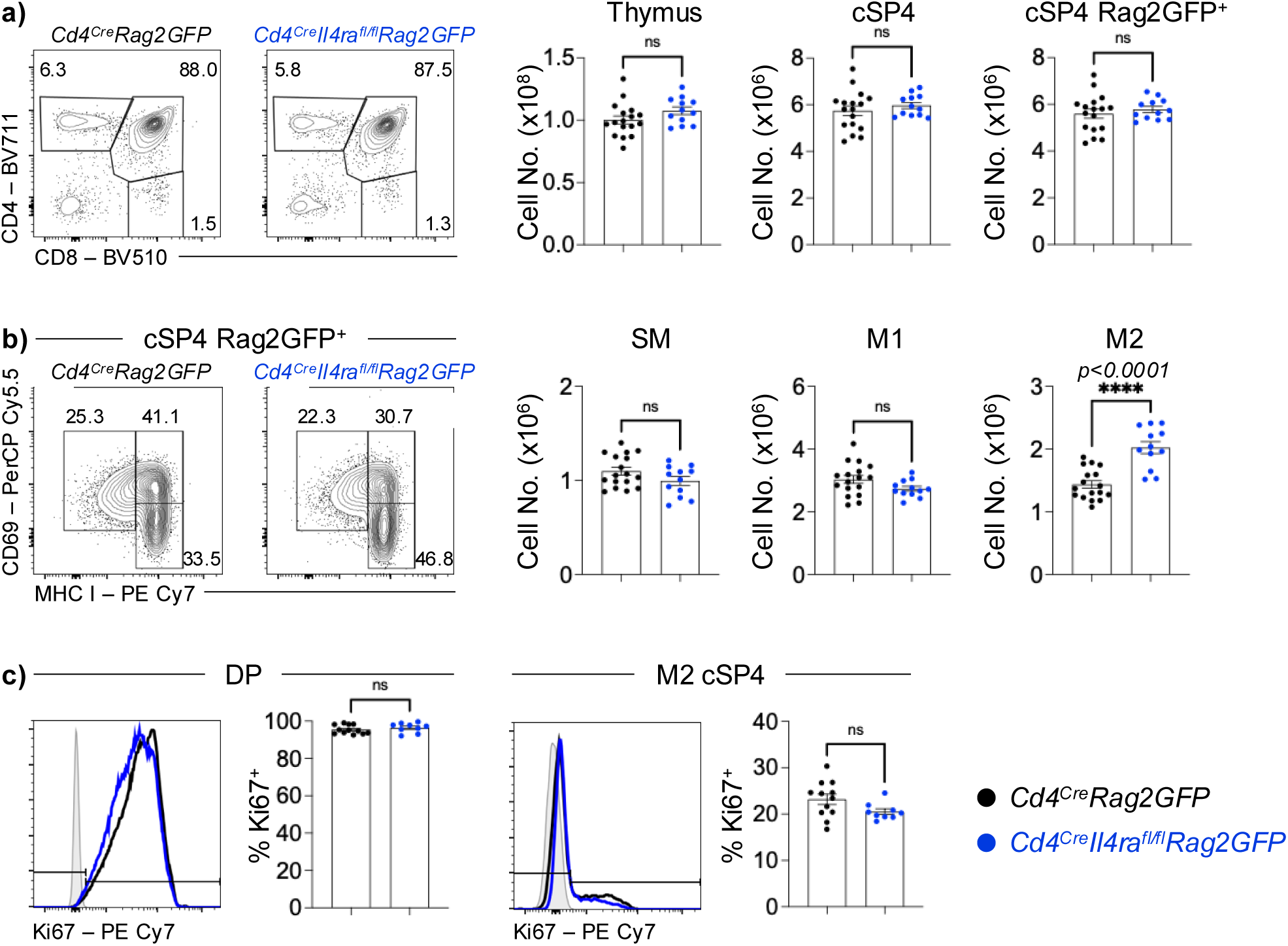
M2 SP Thymocytes Accumulate in the Absence of *Il4rα* Expression. **(A)** Representative flow cytometric analysis of thymocytes from *Cd4^Cre^Rag2GFP* (Black, n=17) and *Cd4^Cre^Il4ra^fl/fl^Rag2GFP* (Blue, n=12) adult mice. cSP4 were gated as PBS57-Tetramer^−^CD4^+^TCRβ^hi^CD25^−^. **(B)** Representative flow cytometric plots, and calculated numbers of SM (CD69^+^MHC I^−^), M1 (CD69^+^MHC I^+^) and M2 (CD69^−^MHC I^+^) subsets of cSP4 Rag2GFP^+^ in *Cd4^Cre^Rag2GFP* (Black, n=17) and *Cd4^Cre^Il4ra^fl/fl^Rag2GFP* (Blue, n=12) mice. **(C)** Representative histogram analysis of Ki67 expression and the frequency of Ki67+ populations within DP (CD4^+^CD8^+^; left) and M2 cSP4 (right) in *Cd4^Cre^Rag2GFP* (Black, n=12) and *Cd4^Cre^Il4ra^fl/fl^Rag2GFP* (Blue, n=9) mice. Data representative of 3 independent experiments.

We have previously shown that cSP4 M2 thymocytes can be further subdivided into a maturational series based on their levels of CD62L, such that M2a cells have the lowest level of CD62L and represent the least mature M2 cells, while M2c express the highest CD62L levels and represent the most mature cells^22^. When we further subdivided total cSP4 M2 cells from *Cd4^Cre^Il4ra^fl/fl^*Rag2GFP mice into M2a, 2b and 2c subsets, we saw that the increase in total M2 cSP4 thymocyes mapped to a specific increase in the most mature M2c cells (Fig. 3A). Similar to cSP4 cells, we also noted an increase in the most mature Rag2GFP^+^TCRβ^hi^ SP8 thymocytes in the thymus of *Cd4^Cre^Il4ra^fl/fl^*Rag2GFP mice (Fig. 2 Supp Fig. 1B-D). Furthermore, while GFP levels in DP, SM and M1 cSP4 subsets from *Cd4^Cre^*Rag2GFP and *Cd4^Cre^Il4ra^fl/fl^*Rag2GFP mice were comparable (Fig. 3B and not shown), we found that both total M2 cSP4 and M2c cSP4 in *Cd4^Cre^Il4ra^fl/fl^*Rag2GFP mice had lower GFP levels than *Cd4^Cre^*Rag2GFP controls (Fig.3 C-D). Interestingly, despite this intrathymic accumulation of mature SP thymocytes, we found that the number of GFP^+^ recent thymic emigrants (RTE) in the spleens of *Cd4^Cre^*Rag2GFP and *Cd4^Cre^Il4ra^fl/fl^*Rag2GFP mice were comparable (Fig. 3E). Additionally, qPCR and flow cytometric analysis showed no change in the levels of the thymus-egress regulator sphingosine-1-phosphate receptor 1 (S1P_1_) in M2 cSP4 thymocytes from *Cd4^Cre^*Rag2GFP and *Cd4^Cre^Il4ra^fl/fl^*Rag2GFP mice (Fig. 3 Supp Fig. 1). Overall, these data demonstrate that IL4Rα deficiency prolongs the length of time mature SP thymocytes spend in the thymus, but does not compromise their ability to exit the thymus and enter into the peripheral T-cell pool.

**Figure. 3:**
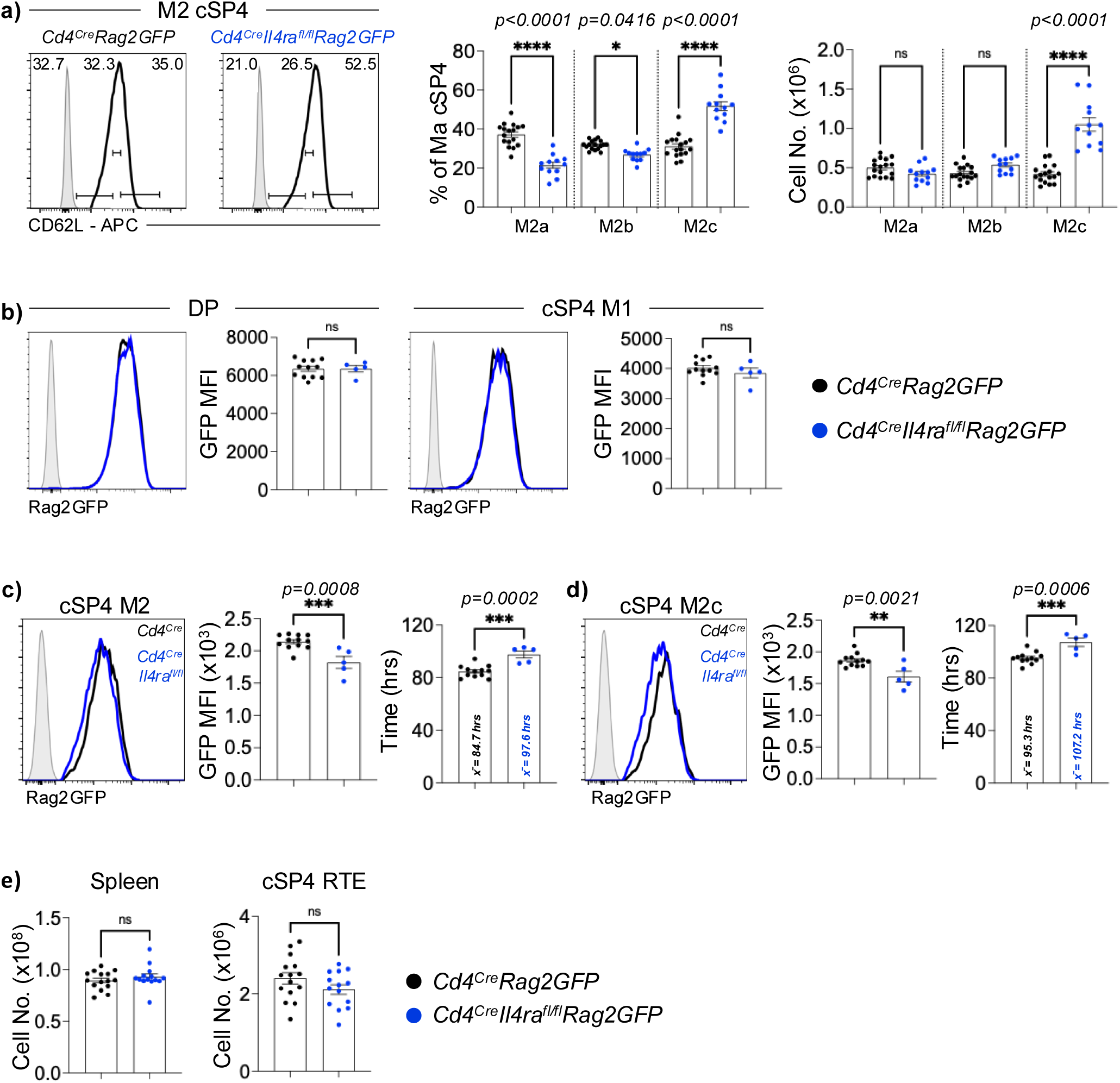
Accumulating M2 SP Thymocytes Are Older in the Absence of *Il4rα* Expression. **(A)** Representative flow cytometric plots, calculated numbers and frequencies of M2a, M2b, and M2c subsets of M2 cSP4 (PBS57-Tetramer^−^ CD4^+^TCRβ^hi^CD25^−^CD69^+^MHC1^+^) in *Cd4^Cre^Rag2GFP* (Black, n=12) and *Cd4^Cre^Il4ra^fl/fl^Rag2GFP* (Blue, n=12) mice. Gating is set by gating M2 cSP4 into 3 equal populations of CD62L^Lo^ M2a, CD62L^Int^ M2b and CD62L^Hi^ M2c in *Cd4^Cre^* control and applying to *Cd4^Cre^Il4ra^fl/fl^*, grey histogram shows isotype staining control. **(B)** Representative histogram of the expression of Rag2GFP by DP (CD4^+^CD8^+^; left), and M1 cSP4 (right) and the plotted mean fluorescent intensity (MFI) of each population in *Cd4^Cre^Rag2GFP* (Black, n=12) and *Cd4^Cre^Il4ra^fl/fl^Rag2GFP* (Blue, n=5) mice. Data from at least 3 independent experiments. **(C + D)** Rag2GFP levels calculated for M2 cSP4 (C) and M2c cSP4 (D) thymocytes from *Cd4^Cre^Rag2GFP* (Black, n=12) and *Cd4^Cre^Il4ra^fl/fl^Rag2GFP* (Blue, n=5) thymus. Grey histograms indicate nonfluorescent control cells. **(E)** Total spleen cellularity and calculated numbers of cSP4 (PBS57-Tetramer^−^CD4^+^TCRβ^hi^CD25^−^) recent thymic emigrants (RTE; Rag2GFP^+^ cSP4) (Black, n=15) and *Cd4^Cre^Il4ra^fl/fl^Rag2GFP* (Blue, n=14) adult mice. For analysis of data in A, a multiple comparison analysis was achieved by an ordinary one-way ANOVA, followed by a Sidak’s multiple comparison posttest in GraphPad Prism to determine statistical differences. Flow cytometric data representative of at least 3 independent experiments. MFI; mean fluorescence intensity.

The experiments outlined above describe the impact of IL4Rα deficiency on SP thymocytes under steady state conditions, where the thymus supports multiple overlapping waves of thymopoiesis in the context of a T-cell replete peripheral immune system. To examine the impact of IL4Rα deficiency under conditions where a synchronised wave of T-cell production and peripheral T-cell pool formation can be monitored from a specific time point, we next established bone marrow chimaeras where congenically marked IL4Rα-deficient bone marrow progenitors were used to reconstitute lethally irradiated WT hosts. In addition, we also utilised a mixed bone marrow chimaera protocol, where equal mixtures of control and gene deficient cells develop alongside each other in the same WT host, allowing us to examine cell-intrinsic versus -extrinsic roles of IL4Rα expression in T-cell development (Fig. 4A). Importantly, compared to mice receiving *CD4^Cre^*bone marrow, mice receiving bone marrow from *CD4^Cre^Il4ra^fl/fl^* mice showed a skewing of cSP4 thymocytes towards the most mature M2c subset (Fig. 4B). Moreover, this skewing of *CD4^Cre^Il4ra^fl/fl^*derived M2 cSP4 thymocytes still occurred in the presence of WT counterparts (Fig. 4B), suggesting that the presence of WT-derived thymocytes does not influence the development of IL4Rα-deficient thymocytes. Finally, when we analysed the peripheral T-cell compartment in the spleen of chimaeric mice at 28 days post-transplant, we saw a reduction in splenic CD4^+^ and CD8^+^ T-cell numbers in chimaeras established from *Cd4^Cre^Il4ra^fl/fl^* compared to controls (Fig. 4C). Collectively, these data demonstrate the requirement for IL4Rα by SP thymocytes is cell intrinsic, and that unlike under steady-state conditions, peripheral T-cell reconstitution following radio-ablation and bone marrow transplantation is influenced by thymocyte expression of IL4Rα.

**Figure. 4:**
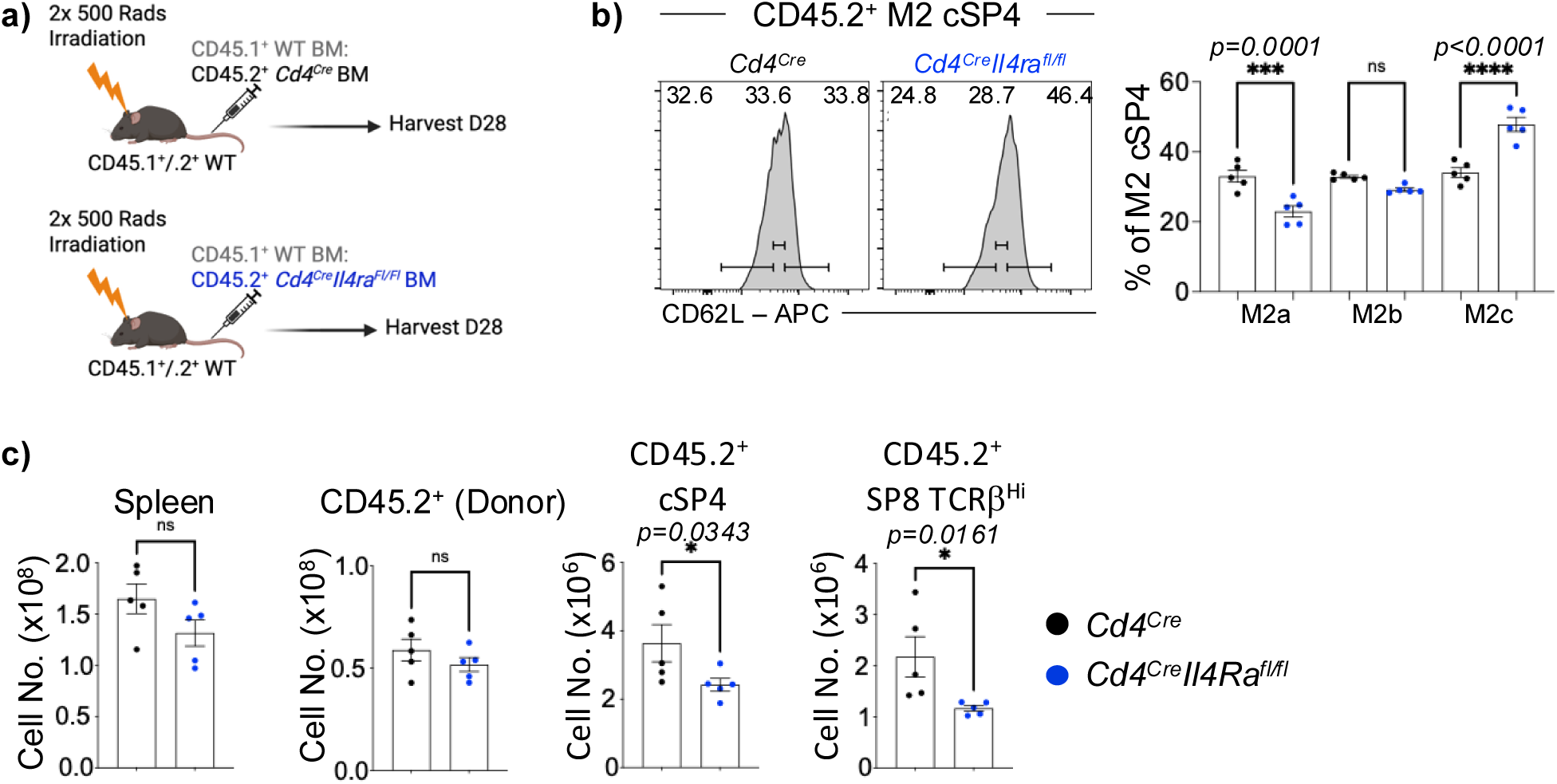
Accumulating M2 SP Thymocytes Are Older in the Absence of *Il4rα* Expression. **(A)** Experimental design for the generation of competitive mixed bone marrow chimeras, where bone marrow from CD45.2^+^ *Cd4^Cre^* or *Cd4^Cre^Il4ra^fl/fl^* was mixed 1:1 with CD45.1^+^ congenically labelled WT bone marrow, and injected into lethally irradiated CD45.1^+^CD45.2^+^ WT host mice. Thymus and spleen were harvested 28 days later for analysis. **(B)** Representative flow cytometric plots, calculated proportions of M2a (CD62L^Lo^), M2b (CD62L^Int^), and M2c (CD62L^Hi^) subsets of M2 cSP4 (PBS57-Tetramer^−^CD4^+^TCRβ^hi^CD25^−^CD69^+^MHC1^+^) in *Cd4^Cre^* (Black, n=5) and *Cd4^Cre^Il4ra^F l/F^* (Blue, n=5) mice. **(C)** Calculated numbers of total spleen cellularity, total CD45.2^+^ donor cells and cSP4 and TCRβ^hi^ SP8 T-cells for *Cd4^Cre^* (Black, n=5) and *Cd4^Cre^Il4ra^fl/fl^* (Blue, n=5) recipients. For analysis of data in B, a multiple comparison analysis was achieved by an ordinary one-way ANOVA, followed by a Sidak’s multiple comparison posttest in GraphPad Prism to determine statistical differences. In all cases, error bars represent mean ± SEM. Flow cytometric data representative of 2 independent experiments.

### Increased Intrathymic Dwell Time Enhances Functional Maturity In Post-Selection Thymocytes

Following thymic selection and prior to thymic exit, the further maturation of SP involves the acquisition of resistance to complement-mediated cell death. This occurs as a result of changes in the incorporation of sialic acid in thymocyte-expressed cell surface glycoproteins. This occurs alongside upregulation of CD55, also known as decay-accelerating factor (DAF), which is a key inhibitor of the complement system^23,24^. This progressive resistance to cell death processes is further accompanied by the acquisition of functional properties, including the ability of M2 cSP thymocytyes to produce cytokines following stimulation^9,24^. At least two models may explain how these processes of continued thymocyte maturation might be linked to thymic occupancy. For example, these changes in behaviour may involve cells reaching a pre-determined intrathymic threshold, meaning that maturity and functional capacities of SP thymocytes are effectively capped once a developmental stage is reached. Alternatively, maturation and functional capacity may be allowed to continue progressively in the thymus. In this scenario, these events would be determined not by developmental stage, but by the length of time SP thymocytes spend in the thymus. To distinguish between these possibilities, we examined whether the increase in intrathymic dwell-time seen in *Cd4^Cre^Il4ra^fl/fl^* mice caused qualitative changes in the behaviour of SP thymocytes.

The incorporation of sialic acids into thymocyte-expressed surface glycoproteins that protects from complement-mediated cell death^24^ can be measured by progressive loss of PNA lectin binding. Thus, PNA binding is greatest at the immature DP stage of thymocyte development, and becomes progressively reduced in maturing SP thymocytes as they incorporate sialic acids^24^, such that low levels of PNA binding indicate increased maturity. In flow cytometric analysis of PNA binding, alongside investigation of levels of CD55 expression, we saw that compared to their counterparts in *Cd4^Cre^* mice, M2 and M2c cSP4 thymocytes from *Cd4^Cre^Il4ra^fl/fl^*showed both reduced PNA binding (Fig. 5A-B) and increased CD55 levels (Fig. 5C-D). Thus, cSP thymocytes from *Cd4^Cre^Il4ra^fl/fl^* demonstrate increased phenotypic maturity, indicating that prolonged thymic occupancy enables the continued maturatrion IL4Rα-deficient SP thymocytes. These findings support a model in which post-selection maturation is not capped at a fixed intrathymic point, but rather suggests it is a continuous process controlled by the period of time SP thymocytes spend in the thymus.

**Figure. 5:**
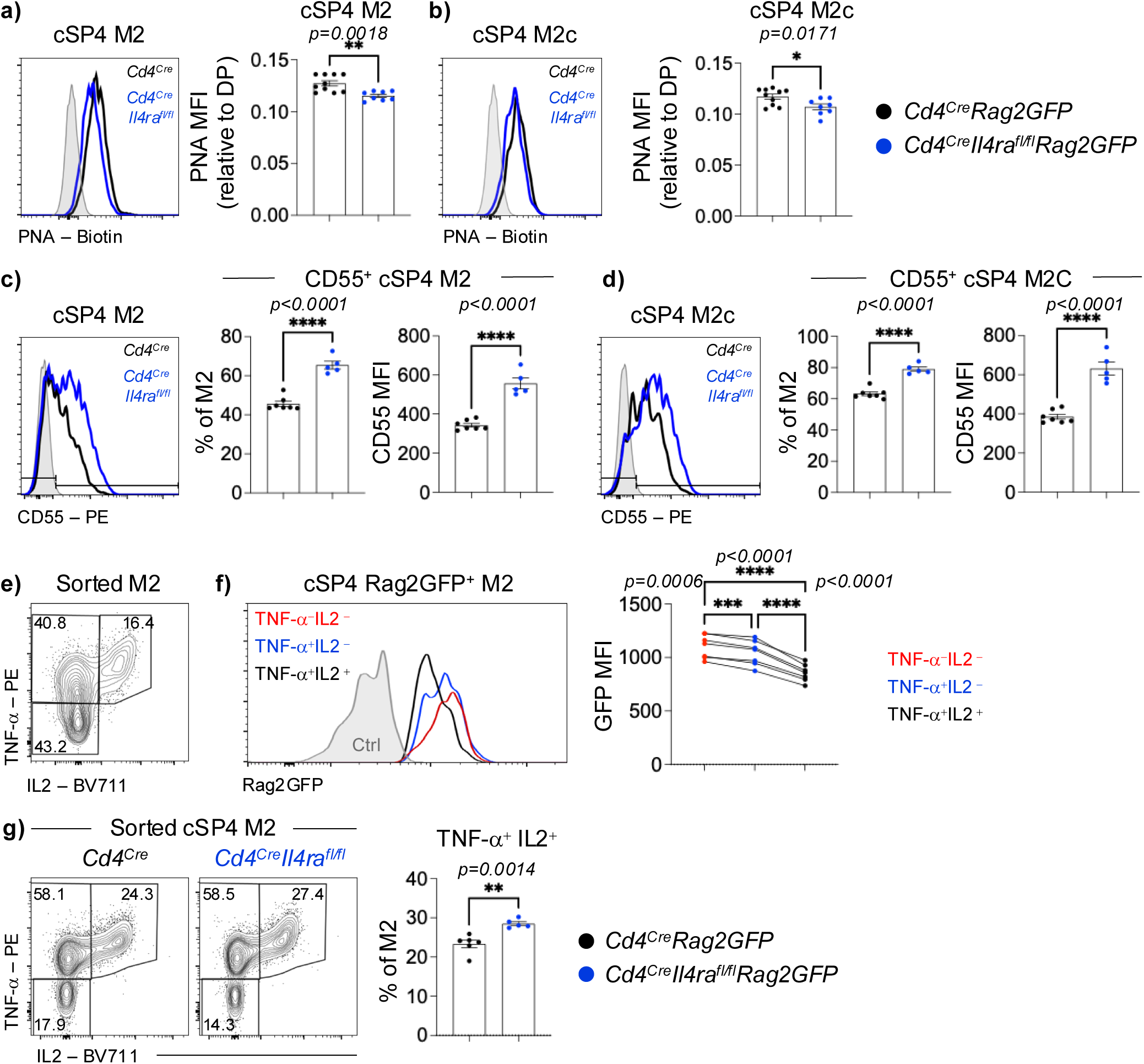
Impact of *Il4ra* Deletion Is Cell Intrinsic. **(A)** Representative histogram of PNA-Biotin expression by cSP4 M2 (PBS57-Tetramer^−^CD4^+^TCRβ^hi^CD25^−^Rag2GFP^+^CD69^−^MHC I^+^) thymocytes from *Cd4^Cre^Rag2GFP* (Black, n=10) and *Cd4^Cre^Il4ra^fl/fl^Rag2GFP* (Blue, n=8) adult mice. Bar chart shows the mean fluorescence intensity (MFI) of PNA-Biotin expression on cSP4 M2, relative to CD4^+^CD8^+^ DP thymocytes. **(B)** Representative histogram of PNA-Biotin expression by the M2c (CD62L^Hi^) subset of M2 cSP4 thymocytes from *Cd4^Cre^Rag2GFP* (Black, n=10) and *Cd4^Cre^Il4ra^fl/fl^Rag2GFP* (Blue, n=8) adult mice. Bar chart shows the MFI of PNA-Biotin expression on cSP4 M2c, relative to CD4^+^CD8^+^ DP thymocytes. **(C)** Representative histogram of CD55 expression by cSP4 M2 thymocytes from *Cd4^Cre^Rag2GFP* (Black, n=7) and *Cd4^Cre^Il4ra^fl/fl^Rag2GFP* (Blue, n=5) adult mice. Left bar chart shows proportion of M2 which are CD55^+^, right bar chart shows the MFI of CD55 expression on CD55^+^ cSP4 M2. **(D)** Representative histogram of CD55 expression by cSP4 M2c thymocytes from *Cd4^Cre^Rag2GFP* (Black, n=7) and *Cd4^Cre^Il4ra^fl/fl^Rag2GFP* (Blue, n=5) adult mice. Bar chart shows proportion of M2C which are CD55^+^, right bar chart shows the MFI of CD55 expression on CD55^+^ cSP4 M2C.**(E)** Representative flow cytometric analysis of TNF-α and IL2 expression by sorted WT adult M2 cSP4 thymocytes following 20-hour anti-CD3/CD28 stimulation + 4-hour PMA/Ionomycin restimulation. N=7 from 2 independent experiments. **(F)** Representative histogram of Rag2GFP expression by M2 cSP4 from WT thymus for TNF-α^−^IL2 ^−^, TNF-α^+^IL2 ^−^, and TNF-α^+^IL2 ^+^ subsets from (E). Rag2GFP MFI is plotted on the right for each subset. A repeated measures ANOVA was carried out, with a Tukey’s multiple comparison post-test. **(G)** Representative flow cytometric analysis of TNF-α and IL2 expression by sorted adult M2 cSP4 thymocytes from *Cd4^Cre^* (Black, n=6) and *Cd4^Cre^Il4ra^fl/fl^* (Blue, n=5) following 20-hour anti-CD3/CD28 stimulation + 4-hour PMA/Ionomycin restimulation. Data from 4 independent experiments.

As cSP4 thymocyte development involves the progressive acquisition of functional properties^9^, we next analysed the process of cytokine licencing in FACS-purified cSP4 M2 thymocytes (Fig. 5 Supp Fig. 1), which produce cytokines in response to TCR stimulation. Interestingly, when we analysed TNFα and IL2 production in conjunction with Rag2GFP levels in sorted and stimulated M2 cSP4 thymocytes, we detected an order in cytokine production that was linked to maturational stage. Here, the least mature (i.e. highest expressers of GFP) subset of cytokine-producing M2 cSP4 thymocytes did not produce cytokine in response to stimulation. In contrast, cells with an intermediate level of GFP produced TNFα but not IL2, while the most mature M2 (i.e. lowest expressers of GFP) produced both TNFα and IL2 (Fig. 5E-F). Importantly, we saw a greater frequency of TNFα^+^IL2^+^ M2 cSP4 thymocytes from *Cd4^Cre^Il4ra^fl/fl^*mice when compared with *Cd4^Cre^* controls (Fig. 5G). Collectively, these demonstrate a developmentally regulated hierarchy in the ability of post-selection SP thymocytes to produce TNFα and IL2. Moreover, they also show that as with maturational status as defined by PNA binding/CD55 levels, functional capacities of SP4 thymocytes are determined by IL4Rα-mediated control of intrathymic dwell time.

### Cellular Diversity Ensures Redundancy In Intrathymic Sources of IL4

The IL4R complex that is expressed by thymocytes consists of a heterodimer formed from IL4Rα and γc chains that binds a single cytokine ligand, IL4^25^. Given the impact of IL4Rα deficiency on multiple aspects of SP thymocytes, we next sought to determine the intrathymic sources of IL4 in the steady-state adult thymus. Specifically, we used the KN2 IL4 reporter mouse strain, where faithful reporting of IL4 protein production is detectable via analysis of human CD2 (HuCD2) expression^26^. Consistent with previous reports^19,27^, production of IL4 by intrathymic cell types was predominantly within iNKT cells, identified using PBS57/CD1d tetamers (Fig. 6A-B). In addition, we also saw a small frequency of SP4 αβTCR^+^ thymocytes and γδTCR^+^ thymocytes that were HuCD2 ^+^ (Fig. 6A-B). As an alternative gating strategy, we also pre-gated on total HuCD2^+^ cells then analysed their subset composition, which again highlighted iNKT-cells as the dominant HuCD2^+^ population in steady state thymus (Fig. 6C). Surprisingly, and despite the well described role of eosinophils as an essential source of IL4 in multiple tissues including the thymus^28,29^, we found that thymic eosinophils were uniformly HuCD2^−^, indicating their lack of active IL4 production (Fig. 6A-C). To reconcile the lack of KN2 expression by eosinophils with their known ability to produce IL4, we next looked directly for the presence of intracellular IL4 protein in thymic eosinophils following permeabilisation and staining with an anti-IL4 antibody. In contrast to other thymic cells, including HuCD2^+^ NKT-cells, the majority of thymic eosinophils were found to be contain intracellular IL4 protein (Fig. 4D-E). Consistent with this, and using an alternative analysis strategy and gating on total anti-IL4^+^ cells, the majority of IL4^+^ cells in the thymus at steady state were eosinophils (Fig. 6F). Taken together, these results indicate that the steady state intrathymic IL4 pool is created by multiple cell types, with iNKT-cells and eosinophils representing the major IL4 producing populations, and that while NKT-cells secrete IL4 following production, eosinophils produce but then retain IL4 intracellularly.

**Figure. 6:**
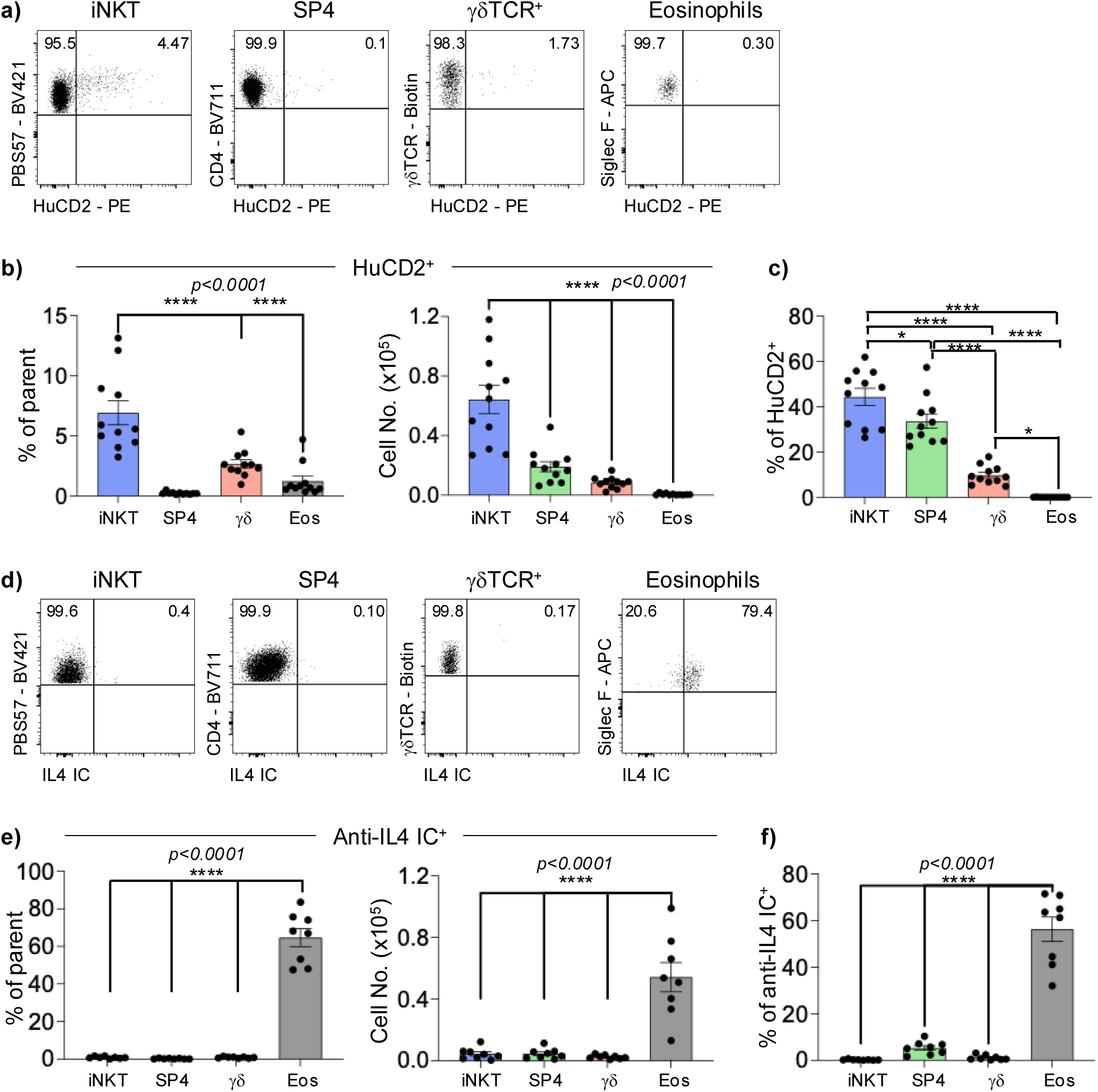
Steady-state IL4 Production Maps To Multiple Intrathymic Cellular Sources. **(A)** Representative flow cytometric analysis of HuCD2 expression in iNKT (PBS57-Tetramer^+^), SP4 (PBS57-Tetramer^−^CD4^+^CD8^−^TCRβ^hi^), γδTCR^+^ (TCRγδ^+^TCRβ^−^) and eosinophils (CD4^+^CD8^−^TCRβ^−^CD11b^+^Siglec-F^+^) from adult KN2 reporter mice. n=11 from 4 independent experiments. **(B)** Quantitation of the proportion (left) and absolute numbers (right) of indicated HuCD2^+^ population within the defined populations from (A). **(C)** Quantitation of the proportion of iNKT, SP4, γδ T-cells and eosinophils within the total KN2^+^ thymus population. **(D)** Representative flow cytometric analysis of intracellular IL4 (IL4 IC) expression in iNKT (PBS57-Tetramer^+^), SP4 (PBS57-Tetramer^−^CD4^+^CD8^−^TCRβ^hi^), γδTCR^+^ (TCRγδ^+^TCRβ^−^) and eosinophils (CD4^+^CD8^−^TCRβ^−^CD11b^+^Siglec-F^+^) from adult WT mouse thymus. n=8 from 4 independent experiments. **(E)** Quantitation of the proportion (left) and absolute numbers (right) of the IL4 IC^+^ population within the defined populations from (C). **(F)** Quantitation of the proportion of iNKT, SP4, γδ T-cells and eosinophils within the total anti-IL4 IC^+^ thymus population. For analysis of data in panel B and E, multiple comparison analysis was achieved by an ordinary one-way ANOVA, followed by a Tukey’s multiple comparison posttest in GraphPad Prism to determine statistical differences. In all cases, error bars represent mean ± SEM. For (C), all comparisons with **** p<0.0001, iNKT vs SP4 p=0.0257 and for γδ vs eos p=0.0468.

To directly test whether iNKT or eosinophils, either individually or in combination, represent essential sources of IL4 for the control of IL4Rα-dependent intrathymic dwell time, we analysed thymocyte development in iNKT-cell deficient (*Cd1d^−/-^*), eosinophil-deficient (ΔdblGATA), and *Cd1d^−/-^*xΔdblGATA double-knockout model (dKO) mice. Here, detailed flow cytometric analysis of late stages of thymocyte development, including proportions and numbers of SM, M1, M2 and M2a/M2b/M2c cSP4 subsets showed that loss of iNKT-cells alone (Fig. 7A), eosinophils alone (Fig. 7B), or both iNKT-cells and eosinophils (Fig. 7C) did not recapitulate the effects on SP thymocytes seen in *Cd4^Cre^Il4ra^fl/fl^* mice. Thus, despite representing the major sources of IL4, iNKT-cells and eosinophils are non-essential regulators of post-selection SP maturation, and points towards cellular redundancy and the involvement of alternative IL4-producing cell types in this process.

**Figure. 7:**
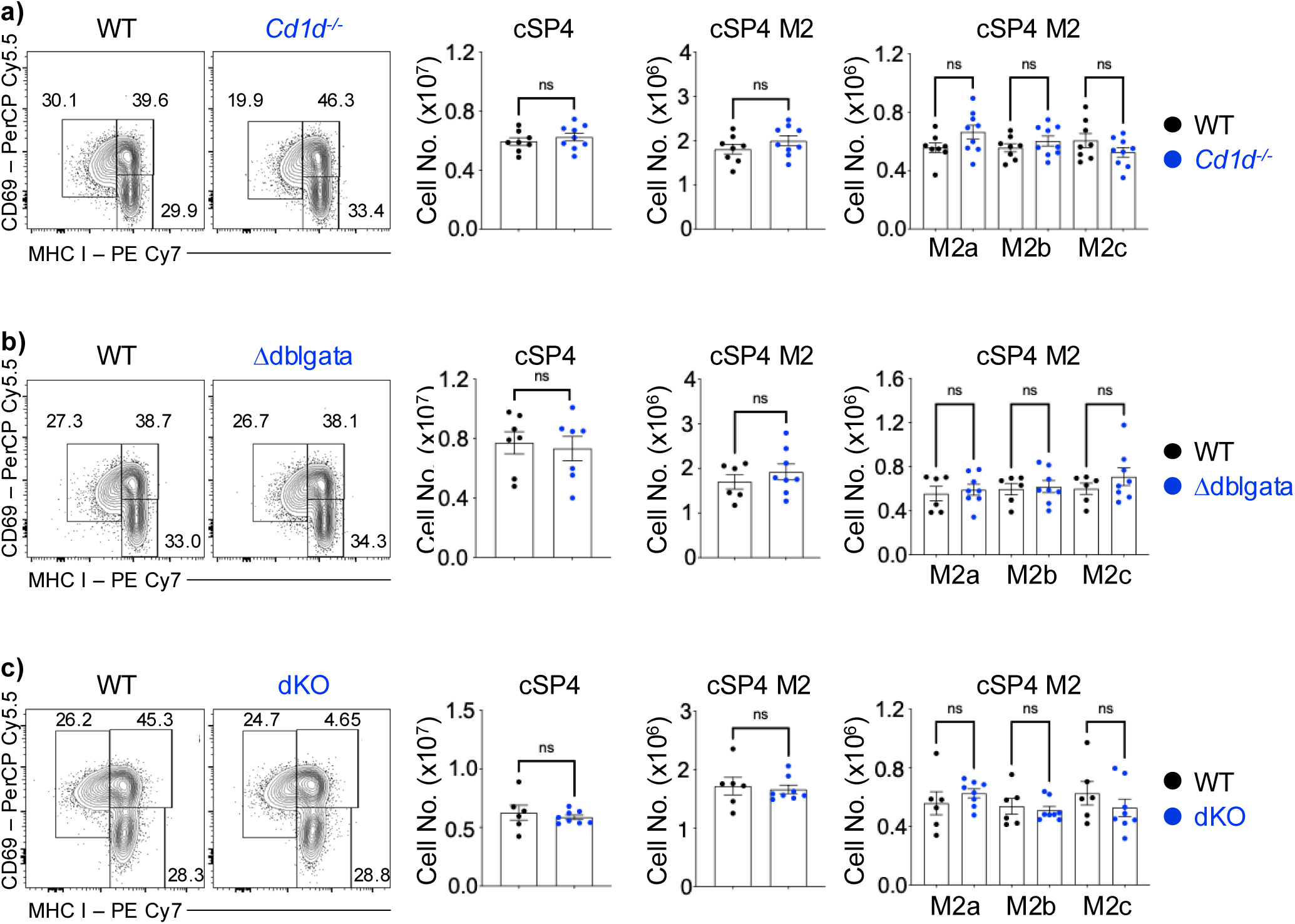
iNKT and Eosinophils Are Dispensable Sources of IL4 In The Control of Thymic Medulla Occupancy. **(A)** Representative flow cytometric plots, and calculated numbers of cSP4 (gated as PBS57-Tetramer^−^CD4^+^TCRβ^hi^CD25^−^), cSP4 M2 (CD69^−^MHC I^+^) and cSP4 M2c (CD62L^Hi^) in WT (Black, n=8) and *Cd1d^−/-^* (Blue, n=9) mice. Data from 3 independent experiments. **(B)** Representative flow cytometric plots, and calculated numbers of cSP4 (gated as PBS57-Tetramer^−^CD4^+^TCRβ^hi^CD25^−^), cSP4 M2 (CD69^−^MHC I^+^) and cSP4 M2c (CD62L^Hi^) in WT (Black, n=6) and Δdblgata (Blue, n=8) mice. Data from 2 independent experiments. **(C)** Representative flow cytometric plots, and calculated numbers of cSP4 (gated as PBS57-etramer^−^CD4^+^TCRβ^hi^CD25^−^), cSP4 M2 (CD69^−^MHC I^+^) and cSP4 M2c (CD62L^Hi^) in WT (Black, n=6) and Δdblgata x *Cd1d^−/-^* double knockout (dKO, blue, n=8) mice. Data from 3 independent experiments. For multiple comparison analysis of M2a/M2b/M2c, an ordinary one-way ANOVA, followed by a Sidak’s multiple comparison posttest in GraphPad Prism was used to determine statistical differences. In all cases, error bars represent mean ± SEM.

### IL4Rα Regulates SP Thymocyte Interactions with mTEC

The findings above indicate that IL4Rα expression determines the length of time SP thymocytes spend in the thymus following their selection. Given the importance of thymocyte:stromal cell interactions for thymocyte development, and the close association of SP thymocytes and medullary thymic epithelial cells (mTEC) in the thymic medulla in both *Cd4^Cre^* and *Cd4^Cre^Il4ra^fl/fl^* mice (Fig. 1), we studied cellular interactions in these mice. In initial experiments, we employed a flow cytometric based tracing system where thymocyte:stromal cell interactions can be quantified through the acquisition by thymocytes of TEC-derived cell membranes that contain MHC class II molecules^30^. Using this approach, we saw readily detectable acquired MHC II on the surface of cSP4 M2 and M2c thymocytes from *Cd4^Cre^* control mice (Fig. 8A). Interestingly, in *Cd4^Cre^Il4ra^fl/fl^*mice, we found that significantly fewer M2 cSP4 and M2c cSP4 thymocytes were MHC II^+^ (Fig. 8A), suggesting IL4Rα-deficient SP thymocytes are impaired in their ability to interact with mTEC.

**Figure. 8:**
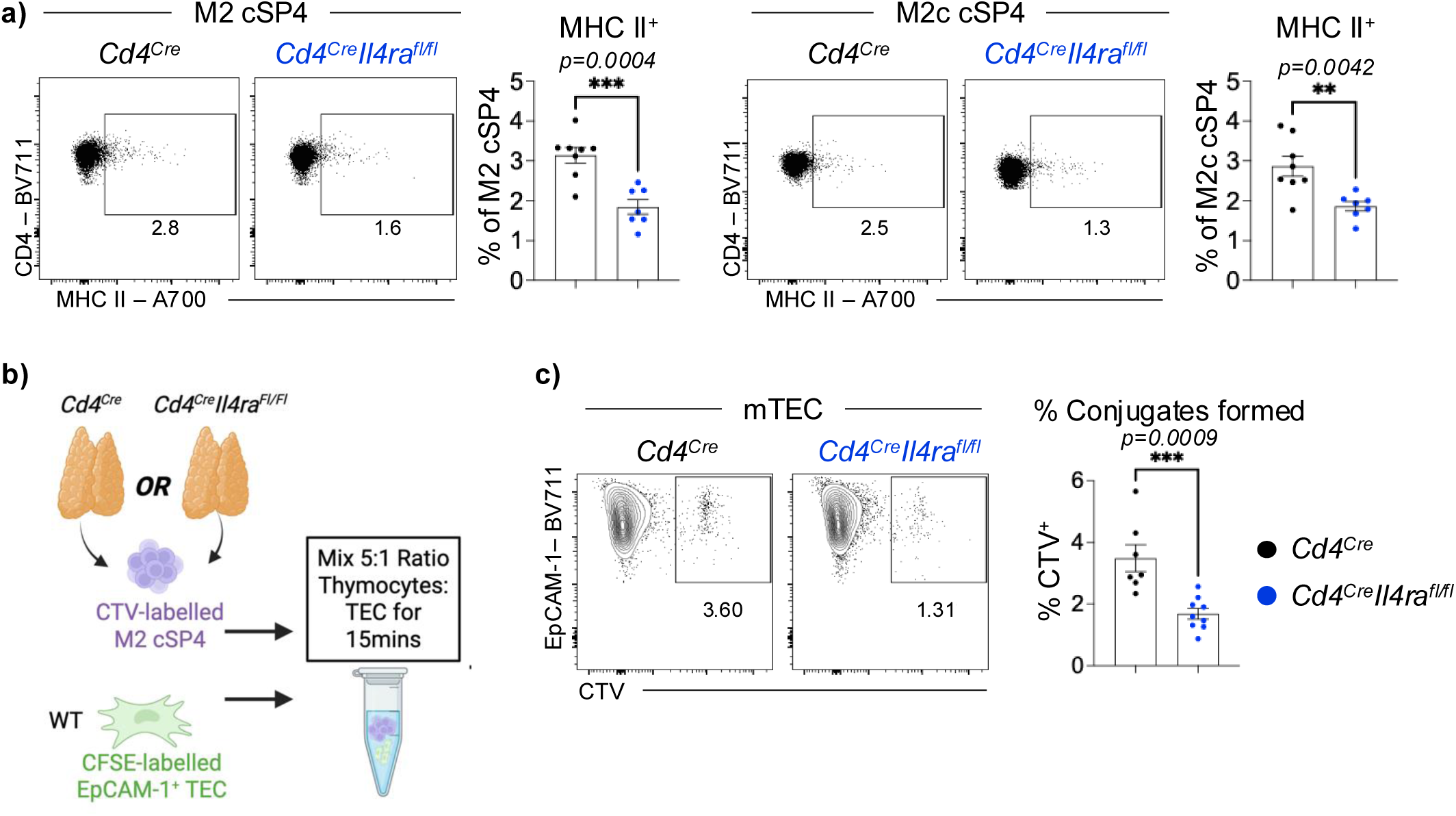
IL4R Controls SP Thymocyte-mTEC Interactions. **(A)** Representative flow cytometric plots and quantitation of the proportions that are MHC II^+^ of M2 cSP4 (gated as PBS57-Tetramer^−^CD4^+^TCRβ^hi^CD25^−^ CD69^−^ MHC I^+^; left) and cSP4 M2c (CD62L^Hi^) (right) from *Cd4^Cre^* (Black, n=8) and *Cd4^Cre^Il4ra^fl/fl^* (Blue, n=7) adult thymus. Plots show the detection of MHC II acquired during their interactions with TEC. Data from 2 independent experiments. **(B)** Experimental approach used to study mTEC:M2 cSP4 thymocyte conjugate formation using flow cytometry. **(C)** Successful mTEC:thymocyte conjugates appear as CFSE^+^EpCAM-1^+^Ly51^−^UEA-1^+^CTV^+^ events within the *Cd4^Cre^* (Black, n=7) and *Cd4^Cre^Il4ra^fl/fl^* (Blue, n=8) mTEC population. Quantitation of mTEC:M2 cSP4 thymocyte conjugate formation is shown on the right. Data from 3 independent experiments.

To directly test this possibility, we next performed flow cytometric-based cell conjugate assays^31^ to quantitate interactions between mTEC and cSP4 M2 thymocytes. FACS-sorted M2 cSP4 thymocytes were labelled with Cell Trace Violet (CTV) fluorescent dye, and then mixed at a 5:1 ratio with EpCAM1^+^ TEC that were FACS-sorted from adult thymus and labelled with CFSE (Fig. 8B; Fig. 8 Supp Fig. 1A-C)^32^. Following 15 minutes incubation to enable conjugate formation, cells were gently resuspended and following gating on Ly51^−^UEA1^+^ mTEC, thymocyte/mTEC conjugates were detected as CTV^+^EpCAM1^+^ events that were quantified by flow cytometry (Fig. 8B). Here, we saw that significantly fewer conjugates (approximate 50% reduction) formed with *Cd4^Cre^Il4ra^fl/fl^* M2 cSP4 thymocytes compared to CD4^Cre^ controls (Fig. 8C). We also saw a smilar reduction in conjugate formation with M2 SP8 thymocytes from *Cd4^Cre^Il4ra^fl/fl^* mice (Fig. 8 Supp Fig. 1D). Collectively, and consistent with reduced MHC II acquisition by SP thymocytes lacking IL4Rα expression, these findings indicate that IL4Rα deficiency in SP thymocytes impairs their ability to interact with MHC II^+^ mTEC within medullary microenvironments, and that failures in successful thymocyte:stromal cell interactions may provide an explanation for the prolongation of their intrathymic occupancy.

## Discussion

αβT-cell production and selection in the thymus is a multi-stage process that guides developing thymocytes through cortical and medullary stromal microenvironments^33,34^. During late stages of thymocyte development that occur following positive and negative selection, a key series of post-selection maturation events enables the thymus to produce functionally competent T-cells that enter the periphery^9^. Despite this importance, the mechanisms that determine post-selection thymic occupancy, and the relationship between thymic occupancy and functional maturity are unclear. By investigating the impact of thymocyte specific loss of IL4Rα expression, we show that lack of IL4Rα selectively prolongs the intrathymic dwell time of the most mature SP thymocytes, and this occurs alongside an increase in their functional maturity prior to thymic exit. Collectively, these results reveal a mechanism in which IL4Rα controls thymic occupancy of SP thymocytes, and shows that this period of thymic occupancy is a key regulator of the maturity of thymic output.

While several studies have analysed the period of time SP thymocytes spend in the thymus^14,15^, and the S1PR1-mediated mechanism that enables thymic exit^11–13^, few studies have examined the triggers that control the duration of thymic occupancy. For example, although intrathymic transfer of immature SP4 thymocytes was shown to increase their maturity, the mechanisms responsible for this were not studied^35^. By defining the importance of IL4Rα in controlling intrathymic dwell time, our data indicates that IL4Rα may act to ensure that SP thymocytes leave the thymus once they have reached a certain stage of intrathymic maturation, ensuring they stay no longer than is necessary. This may then reflect a mechanism to maximise thymic space, and free up niche availability to thymocytes in subsequent waves of development. Such a scenario fits well with previous studies that identified the thymus medulla as an IL4-rich area^17^, and provides an explanation for the presence of this cytokine in this intrathymic site. In addition, that SP thymocytes continue their post-selection maturation during this prolonged period of thymic occupancy indicates that the functional maturity of newly produced T-cells is directly linked to the amount of time they spend in the thymus, rather than SP thymocytes reaching a particular developmental stage.

Importantly, we found that the impact of IL4Rα-mediated control of thymus occupancy on the peripheral T-cell pool depended on whether T-cell development was occurring under steady state or post-transplant reconstitution conditions. Thus, under steady state conditions where multiple waves of thymocyte development and emigration are occurring, prolongued thymus occupancy did not influence the contribution that RTE make to the peripheral T-cell pool. In contrast, during T-cell reconstitution following bone marrow transplantation, prolongued thymic occupancy was accompanied by a reduction in RTE frequency. One possible explanation for this difference is that the phase of peripheral T-cell lymphopenia that occurs following bone marrow transplant may influence thymic emigration and/or the survival of RTE. Further work is required to understand the interplay that takes places between thymic output and the peripheral T-cell compartment at both steady-state and during immune reconstitution. Given the requirement for IL4Rα in post-selection thymocyte development, we also analysed steady-state intrathymic cellular sources of IL4. Here, we utilised the KN2 IL4 reporter strain, which faithfully reports IL4 production, rather than *Il4* gene expression^26^. Consistent with previous studies, iNKT-cells were the most frequent amongst IL4 producing thymic cells ^19,27^, with detectable HuCD2 expression in smaller frequencies of SP αβTCR^+^ and γδTCR^+^ thymocytes, indicating multiple cellular contributions to steady-state IL4 production. Strikingly, eosinophils were found to be HuCD2^−^, which appears at odds with their previously reported role of IL4 production for thymus regeneration^36^. However, direct analysis of the presence of intracellular IL4 protein identified eosinophils as a major intrathymic source of IL4, which is consistent with their ability to store pre-formed cytokines in intracellular vesicles^37–39^. It is interesting to note that IL33 and eotaxin, both of which are known to be produced in the thymus, can trigger eosinophils to release IL4 protein ^38,40–42^. Further studies are required to examine the intrathymic triggers that promote IL4 release from eosinophils in the steady-state thymus. Interestingly, and despite iNKT-cells and eosinophils constituting the majority of thymic IL4-producing cells and residing within medullary areas where they have the potential to influence SP thymocytes^19,43,44^, we found that SP thymocytes were not altered in *Cd1d^−/-^* or ΔdblGATA single mutant mice, or in *Cd1d^−/-^*ΔdblGATA double deficient mice. Thus, IL4 production from iNKT-cells and eosinophils, either individually or in combination, does not explain the requirement for IL4Rα in SP thymocytes. The simplest explanation for this finding is that redundancy occurs in thymic cellular sources of IL4, with αβTCR^+^ SP4 and γδTCR^+^ thymocytes identified here acting as potential additional IL4 providers.

Lastly, our finding that thymocyte/mTEC interactions are influenced by IL4Rα provides an explanation for its importance in controlling the thymic occupancy of SP thymocytes. For example, IL4Rα may control the expression of molecules in SP thymocytes that enable their efficient migration through the epithelial network of the medulla towards thymic exit points. Our finding that cSP4 M2 thymocytes that lack IL4Rα express normal levels of S1PR1 is perhaps consistent with this scenario. Relevant to this, the precise location of exit-niches for thymocytes to leave the medulla and enter the circulation as they undergo post-selection maturation remain unclear. Further studies are required to define the precise intrathymic location of developmentally distinct SP thymocyte subsets within specific medullary microenvironments, which may further uncover mechanisms of intra-medullary thymocyte migration and development.

## Methods

### Mice

The following adult mice of mixed sex and aged between 8 and 12 weeks old were used on a C57BL/6 background,: *Cd4^Cre^* ^45^, *Il4ra^fl^* (originally generated by Frank Brombacher University of Cape Town, Cape Town, South Africa^46^), KN2^26^, Wild type, WT (Charles River), *Cd1d^−/-^* ^47^, ΔdblGATA^48^. The latter two strains were intercrossed to generate *Cd1d^−/-^*ΔdblGATA double knockout mice. Rag2GFP reporter mice^7,49^ were crossed with *Cd4^Cre^* and *Cd4^Cre^Il4ra^fl/fl^* mice. All strains were housed at the Biomedical Services Unit, University of Birmingham.

### Flow Cytometry

Single cell suspensions were generated by mechanical dissociation of thymus lobes and stained with the following antibodies and reagents: anti-CD4 BV711 (RM4-5 BioLegend), anti-CD8 BV510 (53-6.7 BioLegend), anti-CD8 BV785 (53-6.7 BioLegend), anti-TCRβ APC Cy7 (H57.597 eBioscience), anti-TCRγδ Biotin (UC7-13D5 eBioScience), Streptavidin PE Cy7 (eBioscience), Streptavidin BV785 (BioLegend), anti-CD69 PerCP Cy5.5 (H1.2F3 eBioscience), anti-CD62L APC (MEL-14 eBioscience), anti-MHC Class I PE Cy7 (28-14-8 eBioscience), anti-CD25 BV650 (PC61 BioLegend), PBS57-loaded CD1d tetramer (NIH-tetramer facility) BV421, anti-CD55 PE (RIKO-3 BioLegend), PNA biotin (Vector), anti-IL4Rα PE (I015F8 BioLegend), anti-TNFα PE (MP6-XT22 eBioscience), anti-IL2 BV711 (JES6-5H4 BioLegend), anti-Ki67 PE Cy7 (SolA15 eBioscience), anti-S1PR1 (mab7089 R&D Systems), the latter detected with anti-Rat IgG2_A_ Biotin (MRG2a-83 BioLegend). Intracellualr IL4 was detected using anti-IL4 APC (11B11 Invitrogen/eBioscience), cells from KN2 IL4 reporter mice were stained with anti-HuCD2 PE (RPA-2.10 BioLegend). Eosinohils were detected as CD4^−^CD8^−^TCRβ^−^CD11b^+^Siglec-F^+^ cells using the following additional reagents: Anti-SiglecF biotin (Miltenyi) and anti-CD11b BV711 (M1/70 BioLegend). Anti-MHC Class II A700 (M5/114.15.2 eBioscience) was used in the detection of MHC II acquisition by thymocytes from TEC. For detection of thymic epithelial cells, the following reagents were used: anti-EpCAM1 BV711 (G8.8, BioLegend), anti-CD45 APC Cy7 (30-F11 eBioscience), anti-UEA-1 Biotin (Vector Laboratories) and anti-Ly51 APC (6C3 BioLegend).

### pSTAT6 Staining In Thymocytes

Flow cytometric detection of the phosphorylated form of STAT6 (pSTAT6) in thymocytes following in vitro stimulation with recombinant IL4 (rIL4) was performed as previously described^19^. In short, total thymocyte single cell suspensions were incubated in the presence or absence of 10ng/ml rIL4 (Peprotech) for 10 minutes at 37°C, then stained with antibodies to detect the surface markers CD4, CD8, TCRβ, CD69, MHC I and CD62L. Cells were then permeabilised by incubation in 1.6% PFA at 25°C for 10 minutes, followed by methanol at 20°C for 10 minutes and then incubated in anti-pSTAT6 PE (CH12S4N; eBioscience) antibody prior to flow cytometric analysis.

### TNFα/IL2 Cytokine Detection In SP Thymocytes

FACS-sorted M2 cSP4 thymocytes were stimulated in a flat-bottomed 96-well plate with plate-bound 10μg/ml anti-CD3 (145-2C11 eBioscience) and 20μg/ml anti-CD28 (37.51 eBioscience) for 20 hours at 37°C in 5% CO_2_. Cells were washed and restimulated with 50ng/ml PMA (Sigma) and 1.5μM Ionomycin (Sigma) for 4 hours at 37°C in 5% CO_2_, with 10μg/ml Brefeldin-A (Sigma) added after the first hour of incubation. Cells were then washed and stained for surface markers, followed by intracellular cytokine staining using an Intracellular Cytokine Staining Kit as per the manufacturer’s instructions (Becton Dickinson).

### Thymocyte-mTEC Conjugate Assays

Thymocyte suspensions were stained using reagents described above, and M2 cSP4 thymocytes (PBS57-tetramer^−^CD4^+^CD8^−^TCRβ^hi^CD25^−^Rag2GFP^+^CD69^−^MHCI^+^) were sorted using a BD Aria cell sorter (Becton Dickinson). For isolation of EpCAM-1^+^ TEC^36^ adult thymus samples were enzymatically digested using Collagenase Dispase (Roche) and DNase I (Roche). Epithelial cells were enriched by depleting CD45^+^ thymocytes using Miltenyi anti-CD45 microbeads and LS Coloumns, as per the manufacturer’s instructions. Subsequent enriched samples were then stained with anti-EpCAM1 and anti-CD45 antibodies and CD45^−^EpCAM-1^+^ TEC were sorted using a BD Aria cell sorter (Becton Dickinson). FACS-purified thymocytes were labelled with CTV (Invitrogen) while FACS-purified TEC were labelled with CFSE (Invitrogen), as per the manufacturer’s instructions. Following labelling, conjugate analysis was performed as described^31^. Briefly, purified M2 cSP4 and EpCAM-1^+^ TEC were mixed at a 5:1 ratio, centrifuged for 4 minutes at 4°C and 1400 RPM, then gently resuspended in 200μl PBS and incubated at 37°C for 15 minutes to allow conjugates to form. Samples were then gently resuspended and analysed immediately by flow cytometry.

### Immunohistochemistry

Adult thymus tissues were snap-frozen on dry ice and mounted in Tissue-Tek optimum cutting temperature (OCT) compound (Sakura Finetek). Sections were cut at 7µm thickness and fixed in acetone. Sections were then stained using the following reagents: ERTR5 rat IgM (gift from E. Van Vliet, Erasmus University, Rotterdam, Netherlands), detected using anti-rat IgM Alexa Fluor 488 (Invitrogen), anti-CD4 Alex Fluor 647 (RM4-5 eBioscience), anti-CD8 biotin (53–6.7 eBioscience), the latter detected using Streptavidin Alexa Fluor 555 (Invitrogen). Sections were counterstained with DAPI (Invitrogen) to detect nuclei, and confocal microscopy was performed with a Zen 880 microscope (Zeiss), and imaging analysis was performed using Zen Black software (Zeiss).

### qPCR

High-purity complementary DNA (cDNA) was obtained from mRNA labelled with oligo(dT) microbeads using the μMacs One-step cDNA Kit, according to the manufacturer’s instructions (Miltenyi Biotec). Real-time PCR was performed as described^22^ with SYBR Green with primers specific for *Actb* (β-actin) and *S1pr1* on the Corbett Rotor Gene-3000 PCR machine (Qiagen). PCRs were ran in replicates using SensiMix SYBR No ROX Kit (Bioline) with 200 nM of primers for the genes of interest (Sigma-Merck) and *Actb* primers (Qiagen, QT00095242). After an initial polymerase activation step (95 °C for 10 min), cycling was performed for 40 cycles at 95 °C for 15 seconds, 60–62 °C for 20 seconds, and 72 °C for 15 seconds. Amplification specificity was verified by melt-curve analysis. Reaction amplification efficiencies and the Ct values were obtained from the Rotor Gene Real-Time Analysis Software 6.1 using standard curves generated from mouse Universal cDNA Reference-oligo dT primed (BioChain Institute). Fold levels shown in plots represent the mean (±SEM) of replicate reactions and data shown are representative of at least three independently sorted sample sets. Primer sequences used: *S1pr1*: forward 5′-AAATGCCCCAACGGAGACT-3′ and reverse 5′-CTGATTTGCTGCGGCTAAATTC -3′; *Actb* (β-actin): QuantiTect Mm *Actb* 1SG Primer Assay (Qiagen, QT00095242).

### Statistical Analysis

Prism 10 (GraphPad Software) was used to perform all statistical analyses. Unless stated in the figure legend, to compare multiple populations a one-way ANOVA test was used. In all other cases an unpaired Student’s t test was used. In all figures, bar charts and error bars represent means ± SEM, respectively.

## Data Availability

All data generated or analyzed during this study are included in the manuscript and supporting files; source data files have been provided for all figures.

## Acknowledgements

We thank BMSU staff for expert animal husbandry. We thank the NIH Tetramer Core Facility for providing mouse CD1d PBS57-loaded tetramer BV421 reagent.

## Author Contribution Statement

K.D.J., Conceptualization, Data curation, Formal analysis, Investigation, Validation, Writing – original draft.

A.J.W, Data curation, Formal analysis, Investigation, Methodology.

F.M.G., Data curation, Formal analysis, Investigation, Methodology.

S.M.P., Data curation, Formal analysis, Investigation, Methodology.

A.B., Data curation, Formal analysis, Investigation, Methodology.

S.A.P.C., Methodology, Resources.

A.S.M., Methodology, Resources.

A.W., Methodology, Resources.

M.M., Methodology, Resources.

T.L., Methodology, Rescources.

W.E.J., Conceptualization, Project administration, Writing – review and editing.

G.A., Conceptualization, Formal analysis, Supervision, Funding acquisition, Validation, Project administration, Writing – original draft.

---

**Fig. 2 Supp Fig. 1:**
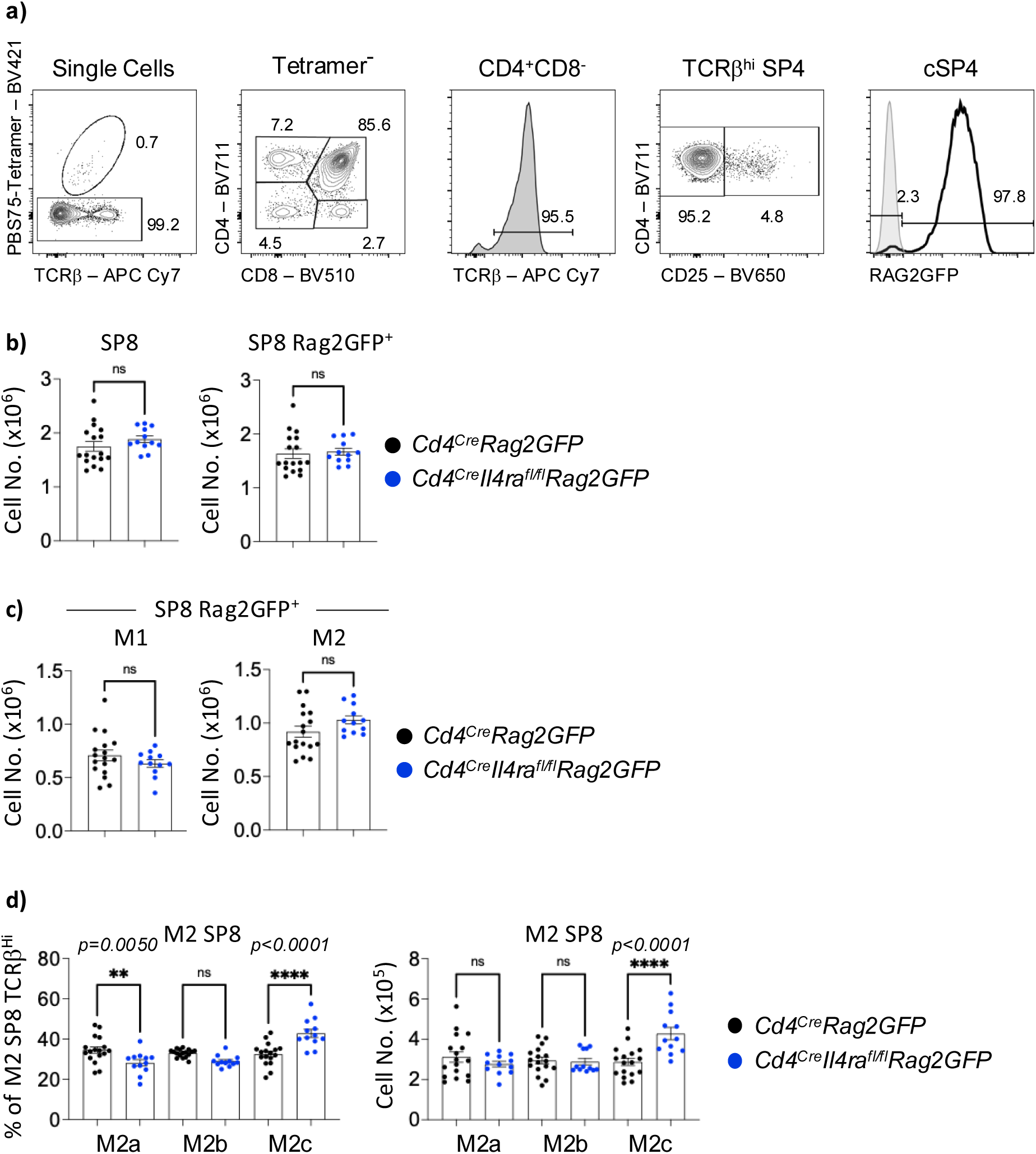
SP8 M2c Accumulate In The Absence of *Il4ra*. **(A)** Gating strategy to identify Rag2GFP^+^ cSP4 thymocytes in *Cd4^Cre^Rag2GFP* thymus. **(B)** Calculated numbers of CD4^+^CD8^−^TCRβ^hi^ SP8 and Rag2GFP^+^ SP8 thymocytes in *Cd4^Cre^Rag2GFP* (Black, n=17) and *Cd4^Cre^Il4ra^fl/fl^Rag2GFP* (Blue, n=12) adult mice. **(C)** Calculated numbers of M1 (CD69^+^MHC I^+^) and M2 (CD69^−^MHC I^+^) subsets of SP8 Rag2GFP^+^ in *Cd4^Cre^Rag2GFP* (Black, n=17) and *Cd4^Cre^Il4ra^fl/fl^Rag2GFP* (Blue, n=12) mice. **(D)** Calculated numbers and frequencies of M2a, M2b, and M2c subsets of M2 SP8 Rag2GFP^+^ thymocytes in *Cd4^Cre^Rag2GFP* (Black, n=17) and *Cd4^Cre^Il4ra^fl/fl^Rag2GFP* (Blue, n=12) mice. For analysis of data in panel C, multiple comparison analysis was achieved by an ordinary one-way ANOVA, followed by a Sidak’s multiple comparison posttest in GraphPad Prism to determine statistical differences. In all cases, error bars represent mean ± SEM. All data collected across at least 3 independent experiments.

**Fig. 3 Supp Fig. 1:**
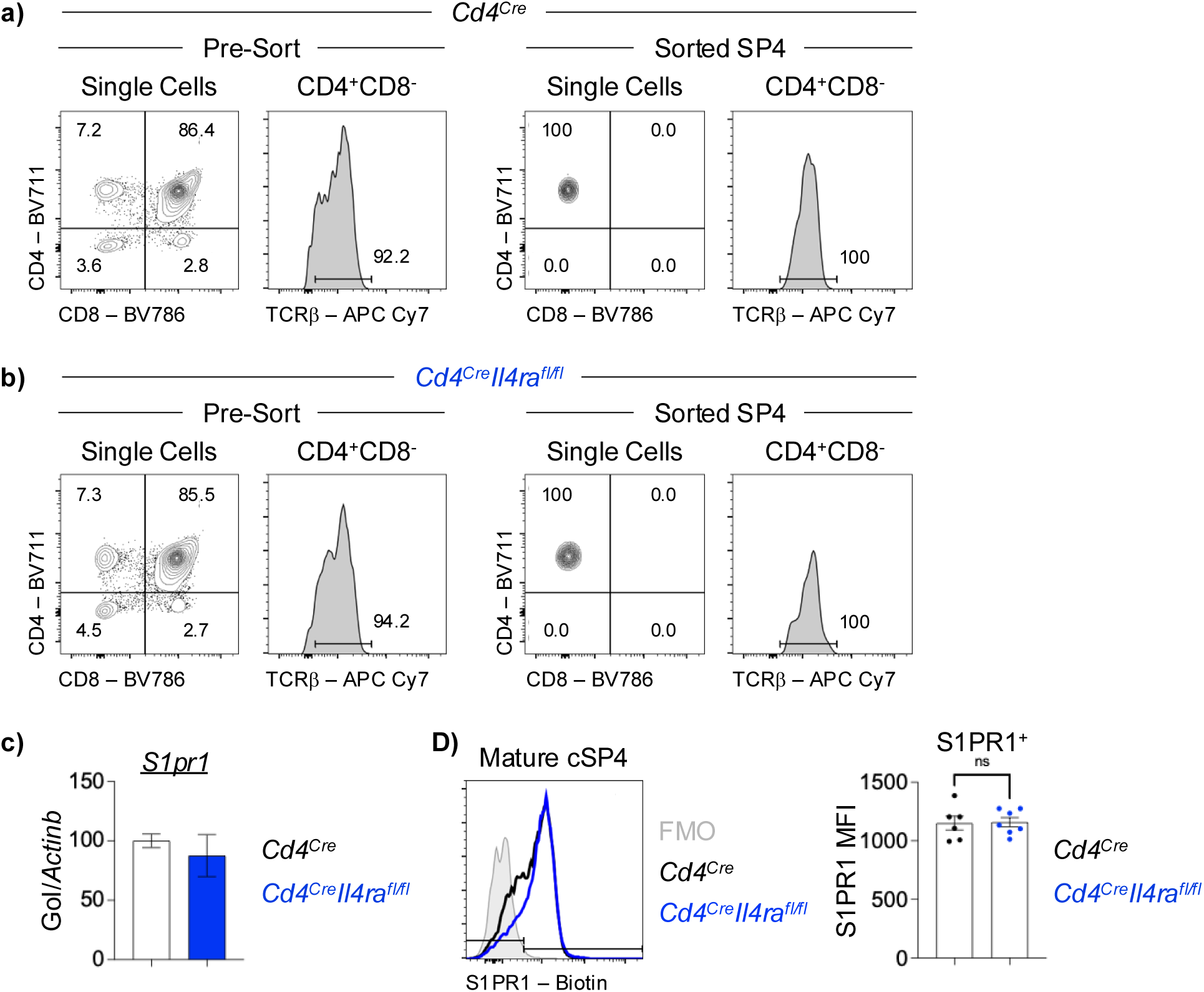
SP Thymocyte Expression Of S1PR1 Is Normal In The Absence of IL4Ra. **(A & B)** Cell sort gating strategy and purity confirmation for the isolation of SP4 thymocytes used in C for PCR analysis from *Cd4^Cre^* (Black) and *Cd4^Cre^Il4ra^fl/fl^* (Blue). **(C)** PCR analysis of *S1pr1* gene expression by sorted SP4 thymocytes from (A + B). Representative of 3 replicates. **(D)** Representative histogram of S1PR1 expression by mature cSP4 thymocytes from *Cd4^Cre^Rag2GFP* (Black, n=6) and *Cd4^Cre^Il4ra^fl/fl^Rag2GFP* (Blue, n=7). The mean fluorescent intensity (MFI) of S1PR1 expression by S1PR1^+^ mature cSP4 is plotted on the right. Grey histogram is an FMO staining control for S1PR1. Data collected across 3 independent experiments.

**Fig. 5 Supp Fig. 1:**
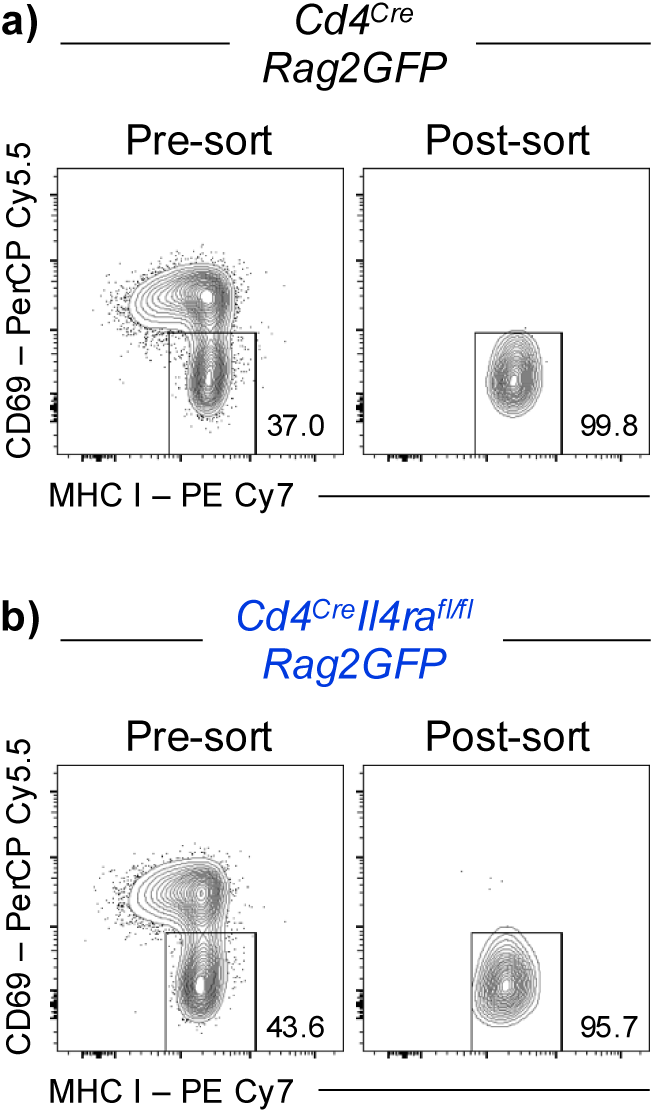
Cell Sort Strategy To Assay Thymocyte Expression of IL2 snd TNFα. **(A & B)** Cell sort gating strategy and purity confirmation for the isolation of M2 cSP4 thymocytes from *Cd4^Cre^Rag2GFP* (A; black) and *Cd4^Cre^Il4ra^fl/fl^Rag2GFP* (B; blue). Representative of 3 independent experiments.

**Fig. 8 Supp Fig. 1:**
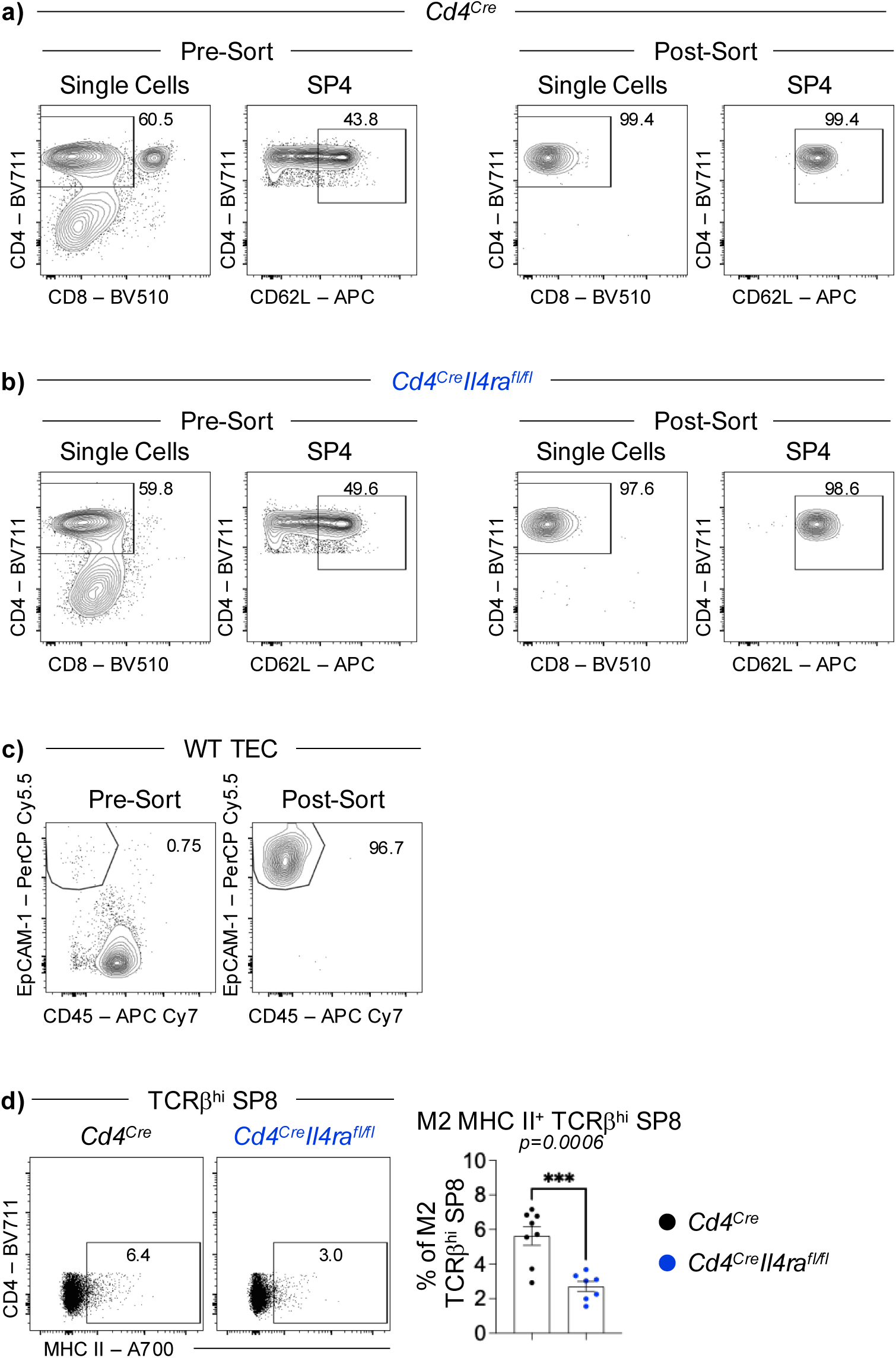
Cell Sort Strategy for Thymocyte-mTEC Conjugate Assays. **(A-C)** Cell sort gating strategy and purity confirmation for the isolation of CD62L^+^ cSP4 thymocytes from *Cd4^Cre^* (A; black) and *Cd4^Cre^Il4ra^fl/fl^* (B; blue). Cell sort gating strategy and purity confirmation for the isolation of EpCAM-1^+^ CD45^−^ thymic epithelial cells (TEC) from WT adult thymus. All data representative of 3 independent experiments. **(D)** Representative flow cytometric plots and quantitation of the proportions of MHC II^+^ M2 SP8 (gated as PBS57-Tetramer^−^CD4^−^ CD8^+^TCRβ^hi^CD69^−^MHC I^+^) from *Cd4^Cre^* (Black, n=8) and *Cd4^Cre^Il4ra^fl/fl^* (Blue, n = 7) adult thymus. Plots show the detection of MHC II acquired during their interactions with TEC. Data from 2 independent experiments.

